# When does the probability of evolutionary rescue increase with the strength of selection?

**DOI:** 10.1101/2024.07.19.604382

**Authors:** Kuangyi Xu, Matthew M. Osmond

## Abstract

Populations may be rescued from extinction via sufficiently rapid adaptive evolution. Evolution is faster with stronger selection but this may come with a demographic cost, creating opposing effects on evolutionary rescue. The outcome of this trade-off determines the optimal strategy for avoiding herbicide/drug resistance evolution. Here we examine the effect of stronger selection, and the associated demographic change, across four models of evolutionary rescue. We find that stronger selection cannot facilitate rescue in two quite different population genetic models unless it increases the absolute fitness of the rescue homozygote. Similarly, in a quantitative genetic model of rescue, where stronger selection leaves maximum fitness unchanged, stronger selection cannot facilitate rescue despite elevating the rate of evolution. We also explore a quantitative genetic model of rescue in a gradually changing environment, generalizing the finding that an intermediate selection strength maximizes survival at steady-state to a wider class of fitness functions.

## Introduction

Populations may avoid extinction via adaptive evolution, a process known as evolutionary rescue (Gomulkiewicz & Holt, 1995). Rescue is of applied relevance in conservation, where it is wanted, and in agriculture and medicine, where it is unwanted. There is now a large literature on evolutionary rescue (reviewed in Bell, 2017; Carlson et al., 2014), including many mathematical models (reviewed in Alexander et al., 2014; Kopp & Matuszewski, 2014). When genotypes do not affect each other’s abundance or fitness, each genotype is an independent population, and rescue depends solely on the initial abundance of genotypes with a positive growth rate (hereafter, rescue genotypes) and their absolute fitness (e.g., rescue via standing genetic variance in haploids; Orr & Unckless, 2008). Consequently, the relative fitness differences between genotypes do not matter (Day et al., 2015). However, genotypes often affect each other’s abundance and fitness, e.g., via mutations (Orr & Unckless, 2008), segregation (Glémin & Ronfort, 2013; Uecker, 2017), and recombination (Uecker & Hermisson, 2016). In this case, rescue can depend on a trade-off between population mean absolute fitness (demography) and the relative fitness advantage of rescue genotypes (selection).

Both demography and selection often co-vary with the severity of environmental change. For example, higher concentrations of herbicides or antibiotics tend to reduce the growth rate of both rescue (resistant) and wildtype (sensitive) genotypes by different extents (Busi et al., 2011; Gould et al., 2018; Gullberg et al., 2011), thus also altering the fitness difference between them. It therefore remains controversial whether higher concentrations of herbicides or antibiotics should lead to a lower likelihood of resistance evolution (Andersson & Hughes, 2012; Bell, 2017; Blanquart, 2019). To illustrate the problem more concretely, consider a locus with alleles *A* and *a* within a diploid population experiencing an abrupt environmental change (e.g., Glémin & Ronfort, 2013; Orr & Unckless, 2008; Uecker, 2017). Assume genotype *aa* cannot grow in the new environment but *AA* can, with *Aa* intermediate (whether the heterozygote can grow or not depends on dominance). The marginal fitness of *A* is dragged down by allele *a*. By eliminating the *a* allele, stronger selection increases the marginal fitness of allele *A* faster, enhancing its establishment probability (Uecker, 2017). However, if stronger selection is associated with a faster decline of *aa* then fewer *A* alleles will arise via mutation during population decline. It is thus unclear how faster population decline and faster evolution combine to affect the rescue probability.

In previous models of evolutionary rescue via large-effect alleles, the strength of selection and the decline rate of the wildtype genotype are typically treated as independent parameters (e.g., Gomulkiewicz & Holt, 1995; Orr & Unckless, 2008; Uecker, 2017). In this case, whether stronger selection increases the probability of rescue depends on how the relative fitness difference is (arbitrarily) assigned to the absolute fitnesses, e.g., decreasing wildtype fitness (Gomulkiewicz & Holt, 1995) vs. increasing mutant fitness (Orr & Unckless, 2008). From a more applied side, there are many pharmacodynamic models that examine the impact of drug dosage on the emergence of drug resistance (reviewed in Blanquart, 2019; Zur Wiesch et al., 2011). Higher drug doses tend to inhibit resistance by removing the sensitive genotypes more rapidly and thereby reducing the number of rescue genotypes that arise by mutation. Higher doses also reduce the growth rate of any given rescue genotype and therefore lower the probability it establishes. While the strength of selection could be measured in this context, if the genotypes do not interact, it is irrelevant (Day et al., 2015).

Incorporating competition between genotypes in models of evolutionary rescue and drug resistance (e.g., Day & Read, 2016; Uecker et al., 2014) implicitly creates an association between absolute fitness and selection. More severe environments accelerate the decline of sensitive genotypes, as in models without competition, but they also reduce competition on resistant genotypes, potentially increasing their absolute fitness and thus selective advantage. This trade-off between the growth rate of resistant and sensitive genotypes can allow the rescue probability to increase, at least to a point, with environmental severity. This is especially true if resistant genotypes are initially abundant since the cumulative input of mutations from the sensitive genotypes (and possibly also the fraction of mutations that can rescue the population; e.g., Anciaux et al., 2018) will decline with environmental severity.

Rescue can also occur via phenotypic selection on quantitative traits (Kopp & Matuszewski, 2014), where the opposing effects of stronger selection also occur (Alexander et al., 2014). Here stronger selection means a larger fitness difference for a given change in phenotype, meaning that a given environmental shift will induce both more rapid evolution and a greater demographic cost. Under an abrupt environmental shift, the larger demographic cost of stronger selection outweighs the benefit from faster evolution (even when holding genetic variance constant) and causes population size to remain below a critical level for a longer period of time (Gomulkiewicz & Holt, 1995), increasing the risk of extinction. The intuition and generality of this result remains unclear. In contrast, under a gradual environmental change with constant variance, stronger selection leads to a lower lag load but a higher variance load (Lande & Shannon, 1996), making persistence most likely at intermediate selection strengths (Lynch & Lande, 1993). This qualitative result remains when stronger selection is allowed to reduce genetic variation (Bürger & Lynch, 1995), which increases the lag load but also decreases the variance load.

The aim of this paper is to identify the conditions under which the probability of rescue increases with the strength of selection, which can cause faster evolution but also pose a greater demographic cost. We consider two genetic architectures, a single locus with two alleles and a quantitative trait. With the single-locus model we examine two cases: 1) rescue genotypes are initially rare and may be lost by chance and 2) rescue genotypes are initially common and extinction risk is driven by stochastic fluctuations in population size. We also consider two scenarios with the quantitative genetic model, abrupt and gradual environmental change.

### Rescue via evolution at a single locus

We first consider a diploid population experiencing an abrupt environmental change. Adaptation involves evolution at a single locus with two alleles *A* and *a*. We consider a discrete-time model and assume the population growth is density-independent. Genotypic absolute fitnesses *W*_*AA*_, *W*_*Aa*_, and *W*_*aa*_ depend only on the magnitude of environmental change *ϵ*. We are interested in environmental changes that cause *W*_*aa*_ < 1 and *W*_*AA*_ > 1, such that a population composed entirely of the *a* allele will initially decline but may avoid extinction if allele *A* reaches a sufficiently high frequency quickly enough.

The strength of selection is described by the selection coefficient, *s* = *W*_*AA*_*/W*_*aa*_ − 1. The fitness of a genotype therefore changes with selection as

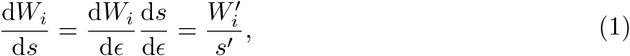

where the prime denotes the derivative with respect to the magnitude of environmental change, *ϵ*. It is also useful to define the dominance coefficient via *hs* = *W*_*Aa*_*/W*_*aa*_ − 1 and the decline rate of the wildtype homozygote, *r* = 1 − *W*_*aa*_.

Denote the frequencies of alleles *A* and *a* at time *t* by *p*_*t*_ and *q*_*t*_ = 1 − *p*_*t*_, respectively. The frequencies of genotypes *AA, Aa*, and *aa* are then 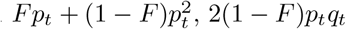 and 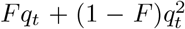, respectively. The inbreeding coefficient, *F*, quantifies the departure of homozygosity from Hardy-Weinberg equilibrium due to non-random mating (Caballero & Hill, 1992). We assume the mating system does not change with the environment such that *F* is constant over time. Particularly, by setting *F* = 1, the model effectively reduces to a haploid population with two genotypes, *A* and *a*. We assume the initial frequency of *A* is small enough that a population of initial size *N*_0_ will suffer demographic decline immediately after environmental change.

### Rescue via the establishment of a rare allele

When allele *A* is initially rare it may be lost by chance. Whether a population will be rescued or not therefore depends on whether allele *A* will establish (Orr & Unckless, 2008). We investigate the probability of rescue either via the establishment of mutations that occur after the environmental change (*de novo* mutations) or via the establishment of *A* alleles that exist at the time of environmental change (standing genetic variation). We assume that mutations from *a* to *A* occur with probability *u* each generations and we ignore mutations back to *a*. For rescue via standing variation, we assume that *A* is selected against before the environmental change, with selection and dominance coefficients *s*\* and *h*\*, and is maintained under mutation-selection-drift balance (Orr & Unckless, 2008).

When allele *A* is rare, its marginal fitness is approximately *W*_*A*_ = *W*_*aa*_(1 + *h*_*e*_*s*). Here *h*_*e*_ = *F* +(1−*F*)*h* is the effective dominance coefficient, describing the fact that when allele *A* is rare it is in a homozygote *AA* with probability *F* and in a heterozygote *Aa* with probability 1 − *F*. We assume allele *A* initially has a positive but small growth rate, 0 *< W*_*A*_ − 1 ≪ 1, so that the establishment probability of an *A* allele is approximately 2(*W*_*A*_ − 1) (Haldane, 1927). The probability of rescue by establishment from *de novo* mutations is then (Glémin & Ronfort, 2013)

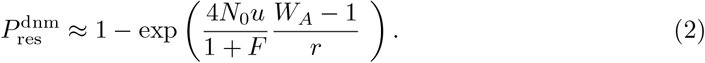

Setting *F* = 1 gives the rescue probability for a haploid population (Orr & Unckless, 2008).

Assuming *N*_0_, *u*, and *F* constant, we see that the probability of rescue from new mutation increases with (*W*_*A*_ − 1)*/r* = *h*_*e*_*s*(1 − *r*)*/r* − 1 and therefore with *h*_*e*_*s*(1 − *r*)*/r* ≈ *h*_*e*_*s/r* and thus ln(*h*_*e*_*s/r*) = ln(*h*_*e*_*s*) − ln(*r*). Taking derivatives with respect to the magnitude of environmental change, *ϵ*, the probability of rescue increases with environmental severity when

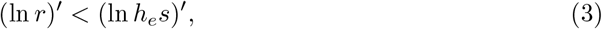

i.e., when the proportional increase in the wildtype decline rate, which reduces mutational input and reduces establishment, is less than the proportional increase in selection on heterozygotes, which increases establishment. It is the proportional changes that matter, mathematically, because rescue depends on the ratio of selection to demographic decline, *h*_*e*_*s/r*. Biologically, stronger selection on the *A* allele, *h*_*e*_*s*, amplifies the detrimental effect of increased decline rate, *r*′ < 0, because that stronger selection is experienced by fewer mutants. Conversely, since the number of mutations decreases like −1*/r*, increasing faster decline rates, *r*, imposes a smaller cost.

Rearranging (3) we see the probability of rescue increases with selection when

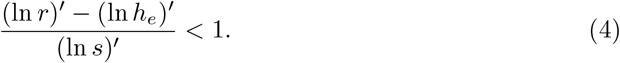

The probability of rescue increases with selection in a larger parameter space when dominance increases with selection, (ln *h*_*e*_)′*/*(ln *s*)′ > 0, as this greatly increases the establishment probability. This is especially true when effective dominance, *h*_*e*_, is lower, as that reduces the cost of fewer mutations. Since (ln *r*)′ ∝ 1*/r* while (ln *h*_*e*_)′ ∝ 1*/h*_*e*_, if *r* ≪ 1, the change in dominance can be ignored and Equation (4) is approximately 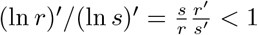, as is with constant *h*_*e*_, and is independent of the inbreeding level, *F*. This condition is always fulfilled when stronger selection is associated with a slower decline of the wildtype homozygote, *r*′*/s*′ < 0 (e.g., lower drug concentrations may increase the fitness of a sensitive strain faster than the fitness of a resistant strain), as we then expect more mutations each with a higher establishment probability. Further, because we are only interested in environments where *W*_*AA*_ > 1 we have *r < s*, which means that the probability of rescue declines with selection when stronger selection is associated with lower fitness of all genotypes, *r*′*/s*′ > 1, as we then expect fewer mutations each with a lower establishment probability.

When stronger selection is associated with a quicker decline of the wildtype homozygote but a higher fitness of rescue homozygote (0 *< r*′*/s*′ < 1), the probability of rescue is more likely to increase with selection when selection is initially weaker and the decline rate of the wildtype homozygote is faster (Figure 1A). As explained below Equation (3), this is because weaker selection reduces the establishment probability of any one mutation and therefore lowers the cost of fewer mutations, i.e., weaker selection makes each mutation worth less to rescue. Similarly, increasing the rate of an already fast wildtype decline has little effect on the expected number of mutations. The same result and reasoning can be reached by considering the marginal costs and benefits of increased selection (Section S1).

**Figure 1:**
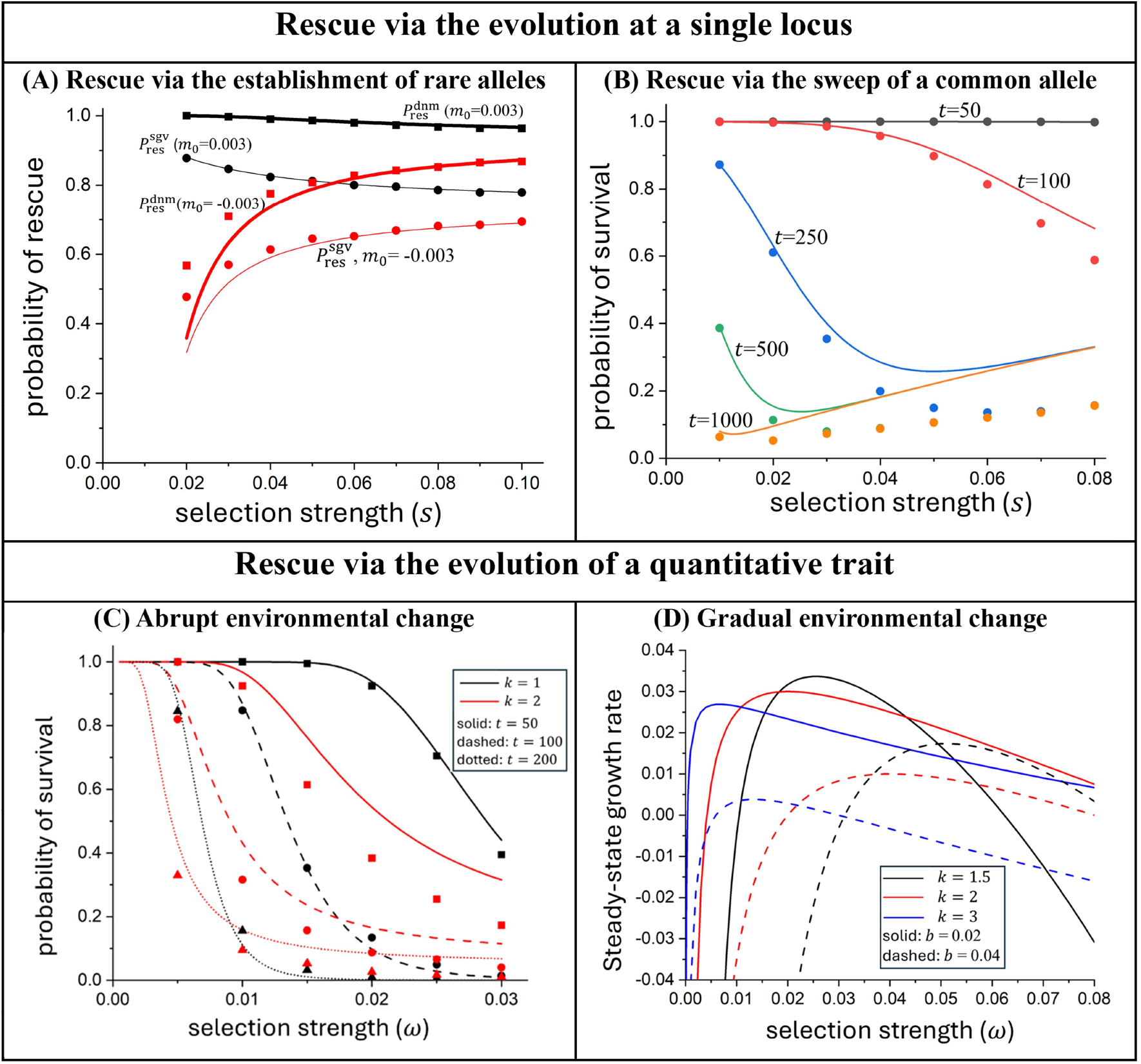
How population survival varies with selection in four different models. Lines are model predictions and dots are from simulations (Section S5). The predictions in **(A)**are from Equations (2) (thick) and (5) (thin). Here (and in panel B) the fitness of each genotype varies linearly with selection: *W*_*aa*_ = 1 + *m*_0_ − *ϕs, W*_*Aa*_ = 1 + *m*_0_ + (*h* − *ϕ*)*s* and *W*_*AA*_ = 1 + *m*_0_ + (1 − *ϕ*)*s*. Parameters values: *ϕ* = 0.3, *h* = 0.5, *F* = 0, *h*\* = *h, s*\* = *s, N*_0_*u* = 1. The predictions in **(B)** are from Equations (S2) and (S6). Parameter values: 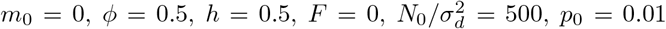. The predictions in **(C)** are from Equations (S2) and (S10), and the first term of Equation (S12). Parameter values: 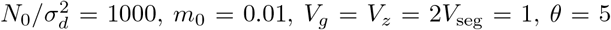. The predictions in **(D)** are from Equation 13. Parameters values: *m*_0_ = 0.05, *V*_*g*_ = *V*_*z*_ = 1.

When rescue is from standing genetic variation, the rescue probability is (Glémin & Ronfort, 2013)

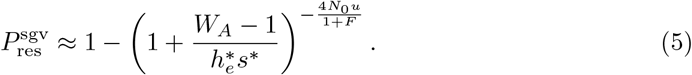

As mentioned in the Introduction, this changes with the severity of environmental change like the absolute fitness of the rescue genotype (Day et al., 2015), *W*_*A*_ − 1 ≈ *h*_*e*_*s* − *r*, i.e., rescue increases with severity when the establishment probability increases, which occurs when selection increases faster than demographic decline. Specifically, 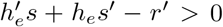. This analysis assumes a fixed number of pre-existing mutant *A*, 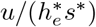, before the environmental change. If we instead imagine that larger-effect mutations are needed to rescue the population from more severe environments, then the number of pre-existing mutants, 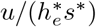, should decline with severity and the probability of rescue will increase with severity when 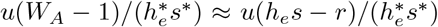 increases. Assuming the effect size of mutants before and after environmental change are proportional, 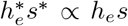, the probability of rescue increases with −*ur/*(*h*_*e*_*s*)). Then in the unlikely case that mutants with different effects are equally likely to arise, *u*′ = 0, by taking the derivative logarithm we get the exact same condition as rescue by new mutation (Equation (3)).

### Rescue via the sweep of a common allele

When allele *A* is initially common, it is almost certain to fix and the change in its frequency is nearly deterministic (Charlesworth, 2020; Hartfield & Glémin, 2016). Under weak selection (*s* ≪ 1), the expected change in allele frequency is approximately (Charlesworth, 2020)

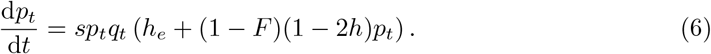

Solving for *p*_*t*_ gives mean population fitness, which in continuous time is

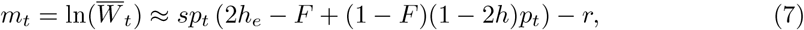

where the approximation assumes weak selection.

With deterministic evolution, population extinction is driven by demographic and environmental stochasticity in population size changes during temporal demographic decline (Xu et al., 2023). Under demographic and environmental stochasticity, the change of population size is, respectively,

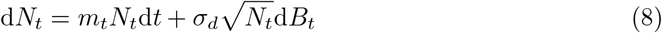

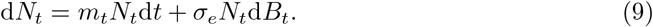

Here d*B*_*t*_ is the Wiener process and 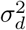 and 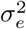 are the strengths of demographic and environmental stochasticity, respectively.

Changes in selection impact the probability a population survives until a time *t* by affecting allele frequency, *p*_*t*_, which influences mean fitness, *m*_*t*_, and determines the expected population size via cumulative fitness, 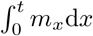. In Section S2 we show that under either demographic or environmental stochasticity, populations have a higher survival probability at time *t* when they have a larger cumulative growth rate for any time *x* ≤ *t* (not *vice versa*).

For the special case of constant additivity, *h* = 1*/*2, in Section S3 we show that the survival probability changes with selection like

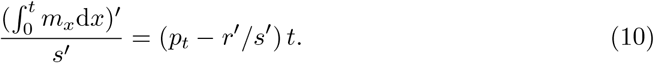

Therefore, the survival probability increases with selection when stronger selection is associated with higher fitness of all genotypes, *r*′*/s*′ ≤ 0, and decreases with selection when stronger selection is associated with lower fitness of all genotypes, *r*′*/s*′ ≥ 1. This latter result indicates that faster evolution is not able to compensate for the cost of a more rapid demographic decline. This is true even in the case of *r*′*/s*′ = 1, such that the maximum fitness, *s* − *r*, is constant. Even then the initial deficit to 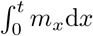 cannot be compensated for by a faster increase in *m*_*t*_, which is perhaps surprising.

Intuitively, in haploids, when increased selection is solely at the cost of a faster decline of the wildtype allele without affecting the rescue allele (*r*′*/s*′ = 1), it will reduce the number of wildtype alleles (*N*_*a*_ = *N*_*a*,0_ exp(−*rt*)) but leave the number of rescue alleles unchanged (*N*_*A*_ = *N*_*A*,0_ exp((*s* − *r*)*t*)), therefore always inhibiting survival. This is also true when the populations is fully inbreeding (*F* = 1). However, when there are heterozygotes increased selection will affect the marginal fitnesses via allele frequency, *p*_*t*_. At early times, stronger selection imposes a greater cost by reducing the marginal fitness of both alleles (Figure 2a-c). However, stronger selection can increase the marginal fitness of both alleles as time proceeds (Figure 2a-c). For example, when *h* = 1*/*2 and *r*′*/s*′ = 1, the marginal growth rate of *A* is *m*_*A*_ = (1 + *p*_*t*_)*s/*2 − *r*, which increases with selection when 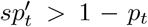. The left-hand side is the benefit of faster evolution and the right-hand side is the cost of reducing heterozygote fitness. The left-hand side peaks mid-sweep, when there is plenty of genetic variance for selection to act on. Evaluating 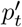 mid-sweep shows that *m*_*A*_ is at least temporarily increased by selection when ln((1 − *p*_0_)*/p*_0_) > 2 (i.e., *p*_0_ < 0.12). Similarly, the marginal growth rate of *a, m*_*a*_ = *p*_*t*_*s/*2 − *r*, can be increased by stronger selection when 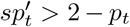, which simplifies to ln((1 − *p*_0_)*/p*_0_) > 6 (i.e., *p*_0_ < 0.0025). Intuitively, stronger selection increases the marginal fitnesses at some time when *A* is sufficiently rare initially, as the benefits of faster evolution then have enough time to accumulate and outweigh the initial cost. The same logic can be applied to dominance; recessive rescue alleles gives more time for the benefits of faster evolution to accumulate (Figure 2a-c). Regardless, increased selection always reduces the expected number of both alleles at any time (Figure 2d-f). This comes down to the fact that the benefit of faster evolution, 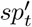, is initially small relative to the cost, before evolution has had much time to change allele frequencies, and after peaking mid-sweep, quickly declines due to a lack of genetic variation at high frequencies.

**Figure 2:**
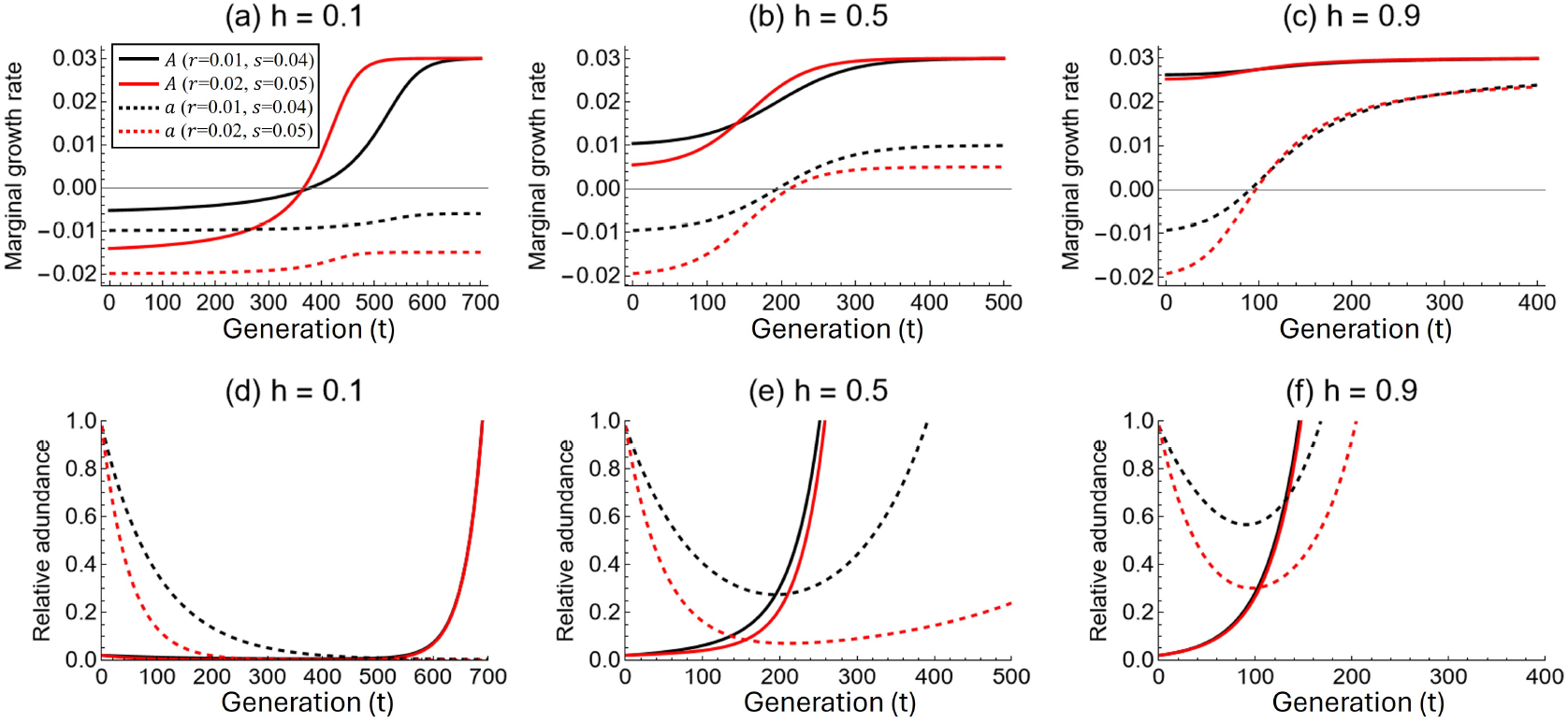
Impact of selection on marginal fitness and abundance of wild-type allele *a* and adaptive allele *A* when rescue occurs via the sweep of a common allele. Increased selection does not change the fitness of *AA*. Parameter values: *p*_0_ = 0.02 and *F* = 0.

When stronger selection is correlated with higher fitness of *AA* and lower fitness of *aa*, 0 *< r*′*/s*′ < 1, cumulative growth rate increases with selection when allele frequency exceeds a critical value (specifically *r*′*/s*′ when *h* = 1*/*2). Therefore, when the initial allele frequency, *p*_0_, is sufficiently low survival declines with selection immediately after the environmental change but potentially increases with selection later (Figure 1B). Because higher effective dominance, *h*_*e*_, increases the rate of evolution and thus allele frequency, *p*_*t*_, survival until a given time is more likely to increase with selection when the rescue allele is more dominant.

### Rescue via the evolution of a quantitative trait

For rescue via the evolution of a quantitative trait, we consider a randomly-mating, sexually- reproducing population in continuous time. The fitness of an individual with phenotype *z* is 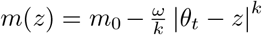, which reaches its highest value, *m*_0_, at the phenotypic optimum, *θ*_*t*_. We assume the phenotype remains normally distributed with mean *z*_*t*_ and variance *V*_*z*_ = *V*_*g*_ + *V*_e_, where *V*_*g*_ is genetic variance, and *V*_e_ is environmental variance.

Assuming the trait distribution is narrow relative to the mean lag behind the optimum, 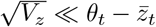, Section S4 shows that the mean phenotype 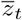 evolves at rate (Lande, 1976)

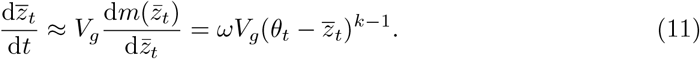

The strength of selection (slope of fitness vs. trait value) depends on both *ω* and *k*. Below we assume a fixed *k* (this is 2 in most previous studies) and refer to *ω* as the selection strength. Like *s* in the population genetic models above, larger *ω* means larger fitness differences between genotypes. Larger *ω* also implies a larger demographic cost (via the second term of *m*(*z*)), as in the population genetic models above when *r*′*/s*′ > 0.

### Abrupt environmental change

We first assume that the phenotypic optimum, *θ*_*t*_, suddenly shifts from 0 to *θ* at time *t* = 0, initiating a demographic decline of a population with initial size *N*_0_. We assume the population is subject to demographic stochasticity so that the dynamics of population size are described by Equation (8). The survival probability depends on the trajectory of mean fitness, *m*_*t*_, which we approximate as 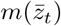, ignoring variance load, and obtain by solving Equation (11).

As shown in Section S2, higher cumulative growth rates imply higher survival probabilities. Assuming constant genetic and phenotypic variance, the cumulative growth rate changes with selection strength *ω* like (see Section S4.1)

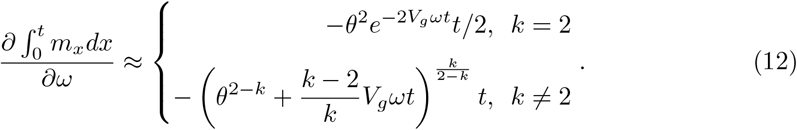

Equation (12) shows that when *k* ≥ 2, stronger selection always reduces the survival probability at any time (Figure 1C) despite hastening evolution. This generalizes the observation at *k* = 2, that the population size at any time is reduced by stronger selection (Gomulkiewicz & Holt, 1995). While this result is known (Alexander et al., 2014), the intuition for why faster evolution cannot make up for a larger demographic cost remains unclear. Considering *k* = 2 for simplicity, the change in mean fitness with increased selection at time *t* is approximately 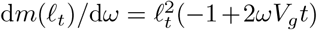, where 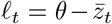 is the mean lag. The consequences of stronger selection scale with the squared lag, which is exponentially declining over time as 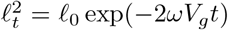 due to evolution. Whether this consequence is a benefit or hindrance depends on the sign of −1 + 2*ωV*_*g*_*t*. The term −1 represents the relative cost of increasing the fitness penalty of a given lag, which is constant. The term 2*ωV*_*g*_*t* represents the relative benefit of a smaller lag through faster evolution, which increases linearly in time as evolution proceeds. Because the relative benefit increases linearly at rate 2*ωV*_*g*_ while the consequence of this benefit declines exponentially at rate 2*ωV*_*g*_, stronger selection cannot increase the cumulative growth rate, 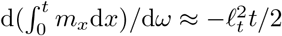 (Equation (12)). In other words, the consequences of stronger selection are larger when the population is more maladapted, which in this case is immediately after the environmental change, before evolution can have much effect. For *k <* 2, Equation (12) only applies for a finite time, 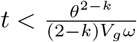, after which the mean phenotype is at the optimum, but stronger selection still tends to reduce the survival probability (Figure 1C).

In simulations, stronger selection reduces the genetic variance and thus the rate of evolution, so that the reduction of survival probability with increased selection is even more severe (Figure 1C). Our calculations above ignore variance load and hence overestimate survival relative to simulations.

### Gradual environmental change

We model a gradual environmental change by assuming the phenotypic optimum increases at a constant rate, *θ*_*t*_ = *bt*. Assuming *k >* 1, the mean phenotype will reach a steady-state lag behind the phenotypic optimum *θ*_*t*_, 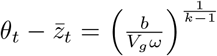 giving a mean fitness of roughly (Section S4.2)

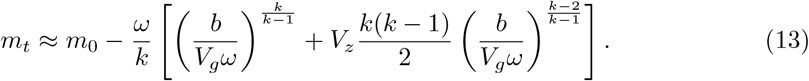

The first term in the square bracket represents the fitness reduction due to the lag of the mean phenotype behind the phenotypic optimum (lag load). The second term is the fitness reduction due to phenotypic variation (variance load), which also depends on the lag of the mean phenotype behind the phenotypic optimum when *k ≠* 2.

Assuming constant genetic and phenotypic variance, the impact of selection strength *ω* on steady-state mean fitness is

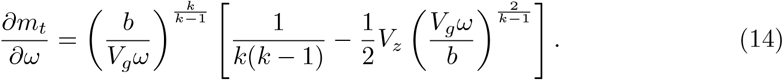

The first term in the square bracket represents the relative benefit of increased selection reducing the lag of the mean phenotype behind the phenotypic optimum. The second term represents the relative cost of stronger selection increasing the variance load.

We see that steady-state fitness, and thus the probability of surviving for a given duration at steady-state (Section S2), is more likely to increase with selection when the environment changes quickly (Figure 1D), the genetic variance is smaller, and the current strength of selection is weaker, as these all increase the lag that can then be reduced by stronger selection. Smaller phenotypic variance reduces the variance load and therefore also allows fitness to increase with selection. The effect of *k* is more subtle, but we see that fitness tends to increase with selection over a larger parameter space when *k* is smaller (Figure 1D). When *k* = 2 the steady-state mean fitness increases with selection when *V*_*z*_ *< b/*(*V*_*g*_*ω*), i.e., when the lag load decreases faster with selection than the variance load increases (Equation (13)).

This implies that an intermediate selection strength is optimal for survival, as previously noted (Alexander et al., 2014), which we generalize to any value of *k* with Equation (14).

Allowing selection to reduce genetic variance yields a very similar steady-state fitness (Section S4.2). This is because reduced genetic variance has opposing effects on fitness, reducing the variance load but increasing the lag load (Figure S1), and these two effects roughly cancel each other out.

## Discussion

Here we have examined how the probability of rescue varies with the strength of selection across four different rescue scenarios (Figure 1) – spanning drug resistance and conservation applications – in an attempt to reach a general consensus. In all models, selection describes the difference in fitness between genotypes, but the approaches differ. In population genetic models of rescue, the selection coefficient is typically treated as an independent parameter that either decreases the absolute fitness of the wildtype (Gomulkiewicz & Holt, 1995) or increases the absolute fitness of rescue genotypes (e.g., Orr & Unckless, 2008). In either case the effect of changing the selection coefficient is clear. However, the strength of selection on a given rescue mutation will covary with absolute fitness. For example, a higher concentration of antibiotics may decrease the absolute fitness of the wildtype faster than the absolute fitness of the rescue mutant, increasing the strength of selection. In this case increased selection comes at a demographic cost, obscuring the consequences on rescue. In quantitative genetic models of rescue, stronger selection – a steeper slope of the phenotype-fitness function – naturally comes with a demographic cost for the concave phenotype-fitness functions typically chosen (e.g., Bürger & Lynch, 1995; Lynch & Lande, 1993). Here, increased selection reduces the absolute fitness of all but the optimal phenotype.

Rescue by rare mutations is determined by the number of mutations that occur and the probability that they establish (Equation (2), Orr and Unckless, 2008, Glémin and Ronfort, 2013). These two factors increase with the absolute fitness of the wild-type homozygote and the heterozygote, respectively. Therefore when stronger selection is associated with lower absolute fitness of both genotypes the rescue probability declines, and vice-versa when stronger selection is associated with higher absolute fitness. The answer is less clear when the absolute fitness of the wildtype homozygote declines with selection but the absolute fitness of the heterozygote increases. Then rescue is more likely to increase with selection when the probability of rescue is low (Equation (4)), as a decline in the absolute fitness of the wildtype homozygote then has a lower marginal cost. This has parallels with the effect of migration on rescue via the establishment of rare alleles (Uecker et al., 2014), as a higher rate of migration increases the number of mutations that experience the new environment but also reduces their selective advantage by exposing them to the old environment more often.

There are some notable differences when switching to a model of rescue via the sweep of a common allele. Here survival increases with the cumulative mean growth rate, which in turn increases with the absolute fitnesses of the genotypes and the allele frequency (Equation (7)). Despite allele frequency increasing faster with stronger selection, stronger selection still cannot help rescue if it does not increase the fitness of some genotypes, at least under additivity (Equation (8)). This is surprising given that in this case stronger selection can increase mean fitness at a given time. It turns out that this benefit of faster evolution is unable to compensate for the demographic cost of lower mean fitness immediately following the environmental change. Increased selection can help rescue when it is associated with increased fitness of the mutant homozygote. Survival is then increased when allele frequency is above a critical value, which occurs earlier when selection is more efficient (higher dominance coefficient) and initially stronger. This latter result is the converse of the result for rescue by new mutation, where rescue is more likely to increase with selection when the rescue allele is recessive and selection is initially weak. With a common allele, selection influences survival via *sp*_*t*_ − *r*, rather than via *s/r*, and therefore initially stronger selection, *s*, translates faster evolution, 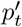, into larger fitness gains. This has important implications because increasing drug doses may enhance the fitness of resistant strains, for example, by reducing competition from susceptibles (Day & Read, 2016; Day et al., 2015; Huijben et al., 2011; Lipsitch & Levin, 1997; Uecker et al., 2014). Whether higher drug doses increase rescue via competitive release therefore strongly depends on the mode of rescue.

The quantitative genetic model of rescue following an abrupt environmental change gives decidedly different results than any other model we considered. In particular, stronger selection always reduces survival (Equation (12)). This was already known for a quadratic (continuous-time) fitness function (Alexander et al., 2014), but here we generalize this to fitness functions of the form *m*(*z*) = *m*_0_ − (*ω/k*)|*θ*_*t*_ −*z*|^*k*^ for any *k* ≥ 1. As in the population genetic model of rescue via the sweep of a common allele, the benefit of faster evolution and even higher growth rates at later times cannot compensate for the demographic cost immediately after the environmental change. It is curious that these two models coincide on this point despite their different functional forms. Determining what needs to be true for faster evolution to make up for a larger initial demographic cost remains an open question.

The quantitative genetic model with a gradual environmental change is distinct from the others because with a gradual change we examine the population at a steady-state, ignoring the transient dynamics. Here it is well-known that an intermediate strength of selection is optimal for survival at steady-state (Alexander et al., 2014). Stronger selection reduces the lag load via faster evolution (like it can increase mean fitness in the common allele and quantitative genetic models of an abrupt change above), but it also increases the variance load. Stronger selection therefore helps survival when the lag is initially large and the phenotypic variance small. Here we generalize this result to fitness functions of the form *m*(*z*) = *m*_0_ − (*ω/k*)|*θ*_*t*_ − *z*|^*k*^ for any *k* ≥ 1 (Equation (14)). Stronger selection tends to increase survival over a larger parameter space when *k* is smaller as the variance load is then relatively smaller (Equation (13)). With *k <* 1 the selection gradient becomes weaker as lag increases, leading to instability (see also Osmond & Klausmeier, 2017).

One could extend this investigation to ask how selection affects rescue in more complex models, considering, for example, alleles at multiple loci (Gomulkiewicz et al., 2010; Uecker & Hermisson, 2016) responding to drug combinations or multiple life-stages (Barfield et al., 2011; Cotto et al., 2019), where the rate of evolution and demography feedback. One particularly exciting direction would be to model competition between genotypes (Day & Read, 2016; Uecker et al., 2014) as this is a scenario where the optimal severity of environmental change for survival (or extinction) is uncertain. Because competition is a mechanism by which the decline rate of the wildtype is positively associated with a larger selective advantage of the mutant (over some environmental severities), we suspect this model might behave similarly to the population genetic models explored here with *r*′*/s*′ > 0. Analogously, it would be interesting to consider the effect of selection on rescue when there is competition between species. For example, an increase in herbicide dose can alter selection in different species to varying degrees, thereby changing their relative competitiveness, which in turn affects resistance evolution in each species.

## Data availability

The simulation code is attached as Supplementary Files.

## Supplementary materials

### S1 Rescue via the establishment of a rare allele

To better understand the roles of *r* and *s* in the cost and benefit of increased selection on rescue by new mutations, note that the probability of rescue increases with the product of the total number of *A* alleles that arise, *n*_*A*_ = 2*N*_0_*u/*((1 + *F*)*r*), and the establishment probability of each, *π* = 2(*W*_*A*_ − 1) ≈ *h*_*e*_*s* − *r*. The marginal effect of increased selection on this product is

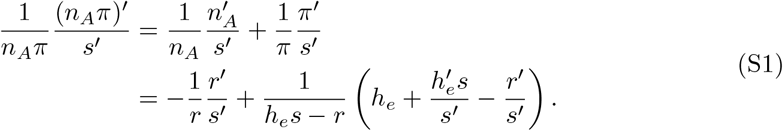

Assuming *r*′*/s*′ > 0, the first term is the relative cost of reduced mutational input. Assuming 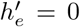 for simplicity, the second term is the relative benefit of higher establishment probability. The probability of rescue increases with selection when the marginal benefit is greater than the marginal cost, which reduces to Equation (4).

### S2 A sufficient condition for a higher population survival probability under demographic or environmental stochasticity

Here we prove a sufficient condition for a higher population survival probability when the population size change is subject to demographic or environmental stochasticity. Consider two mean fitness trajectories over time, *m*_*t*_ and 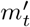. Denote the survival probability under *m*_*t*_ and 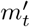 by *P*_surv_(*t*) and 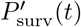, respectively. We show that if 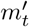 gives a higher cumulative growth rate than *m*_t_ up to time *t*, i.e., 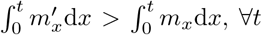, then under either demographic or environmental stochasticity, the survival probability is higher, 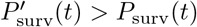.

Under demographic stochasticity we consider a population extinct when *N*_*t*_ = 0. The probability that a population is not extinct at time *t* is then (Anciaux et al., 2018)

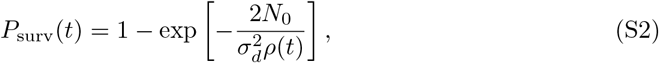

where 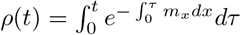. Since 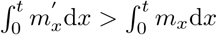 at all *t* then 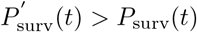 at all *t*.

Under environmental stochasticity Equation (9) indicates that the population size at time *t* is

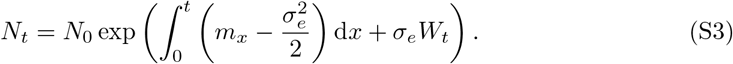

Since *N*_*t*_ is always positive, we assume that the population becomes extinct when *N*_*t*_ is drops below a threshold *N*_*c*_. Analytical solutions to the survival probability are not available when *m*_*t*_ changes over time. Instead, let the time that the population becomes extinct be *t*\* = inf {*t >* 0|*N*_*t*_ ≤ *N*_*c*_} so that the survival probability is *P*_surv_(*t*) = 1 − *P* (*t*\* ≤ *t*). Then, since 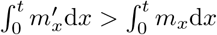 for all *t* we have

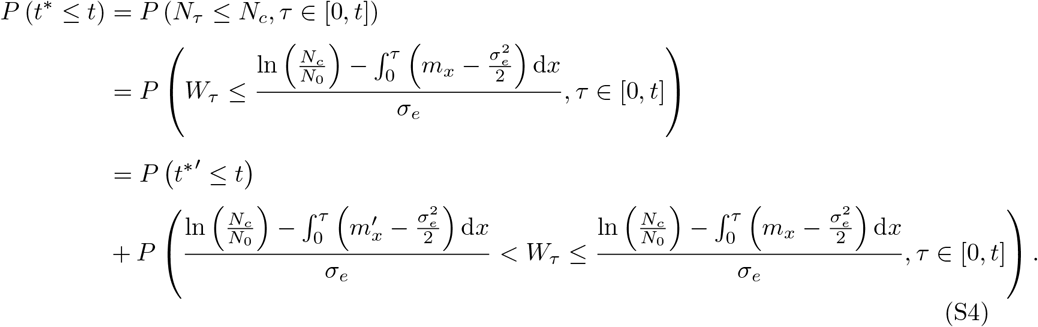

Therefore *P* (*t*^*\*′*^ ≤ *t < P* (*t*\* ≤ *t*), so that 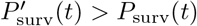 at all *t*. Intuitively, since the expected population size at time *t* is larger with a larger cumulative growth rate up to time *t*, the probability of extinction is smaller.

### S3 Rescue via the sweep of a common allele

The change in frequency of a common rescue allele under weak selection is given by Equation (6). For the special case of constant additivity, *h* = 1*/*2, we can solve for the allele frequency at time *t*,

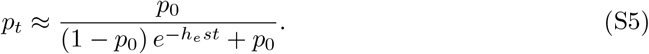

From Equation (7), under constant additivity the cumulative growth rate is

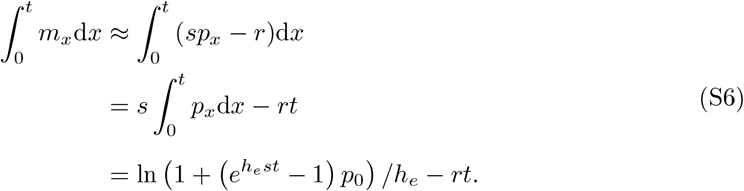

This changes with selection like

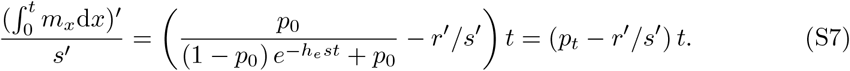

### S4 Rescue via the evolution of a quantitative trait

Denote the genetic value of an individual with phenotype *z* by *g*. Based on the Price equation (Price, 1970), the rate at which the mean phenotype evolves is

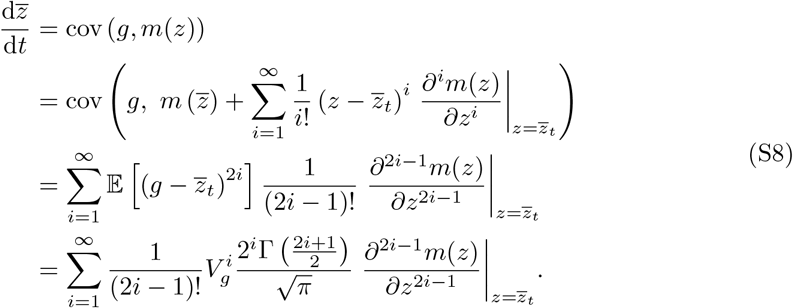

Since 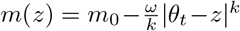 we have 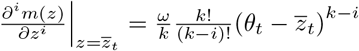 for 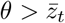 Therefore, if the distribution of genetic values, *g*, concentrates sharply around the mean phenotype, 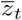, compared to the environmental shift, 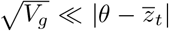, we can approximate the rate of change in the mean trait value with just the leading term,

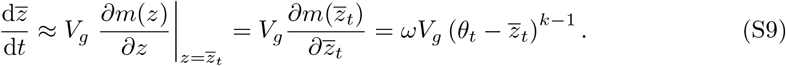

See Iwasa et al., 1991 for a similar derivation in discrete time.

#### S4.1 Abrupt environmental change

Under an abrupt environmental change from *θ*_*t*_ = 0 to *θ*_*t*_ = *θ* at time *t* = 0, from Equation (S9) the mean trait value at time *t* is

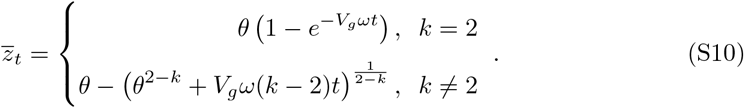

For *k* = 2 the solution is exact. For *k >* 2, the approximation becomes inaccurate as *t* → ∞ because 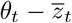 becomes the same order as *ωV*_*g*_, violating the implicit assumption leading to Equation (S9). When *k <* 2, the approximation is only accurate until 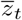 evolves to *θ*, at 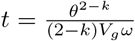.

To see how the strength of selection affects the cumulative growth rate we need to calculate 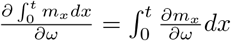. The mean population fitness can be expanded as above,

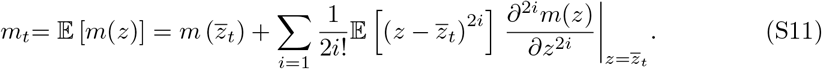

Assuming a narrow trait distribution relative to the optimum shift, 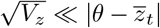, we can use just the first two terms,

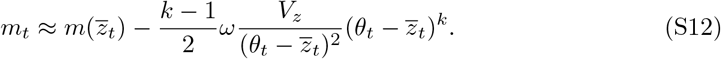

The first term declines with the mismatch between the mean trait value and the optimum (maximum fitness minus lag load) and the second term becomes more negative with phenotypic variation (variance load).

Ignoring the second order term, Equation (S12) becomes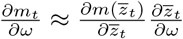, which means that changes in selection strength *ω* affect mean growth rate at time *t* via the rate of decay of fitness around the optimum, 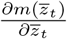, and the rate of evolution of the mean phenotype, 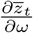. The change in mean growth rate with selection is then

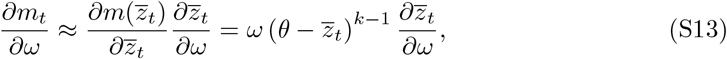

where the change in mean trait with selection, from Equation (S10), is

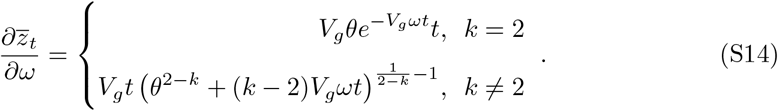

Equation (12) is obtained by substituting Equations (S13) and (S14) into 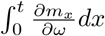.

#### S4.2 Gradual environmental change

Under gradual environmental change, we assume the phenotypic optimum changes at a constant rate *b* over time, *θ*_*t*_ = *bt*. The mean, 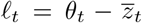, changes like 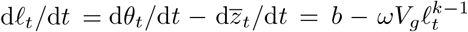, giving steady-state 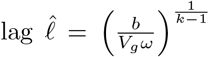. The eigenvalue (d(d*𝓁*_*t*_*/*d*t*)*/*d*𝓁*_*t*_) evaluated at the steady state is 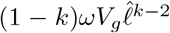, meaning the steady-sate lag is stable when *k >* 1 and unstable when *k <* 1 (because selection gets weaker, and hence evolution slower, as the lag grows). The mean growth rate at the steady state can be obtained by plugging the steady-state into Equation (S12), which gives Equation (13).

In the main text, we analyze the effects of selection strength on the steady-state growth rate by assuming constant genetic variance across different selection strengths. However, in reality, genetic variance will change as selection strength changes, since genetic variance is determined by a balance between the introduction of new variation (e.g., segregation, mutation) and the removal of variation by selection. To incorporate the reduction of genetic variance with increased selection, we adopt the infinitesimal model (Barton et al., 2017). Briefly, the model assumes that the genetic value of an offspring is

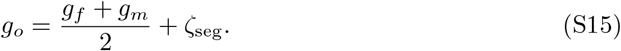

The first term is the expected genetic value, which is the average genetic value of its mother (*g*_*f*_) and father (*g*_*m*_). The second term, ζ_seg_, is the noise due to genetic processes such as segregation and recombination, which is assumed to be normally distributed with mean 0 and variance *V*_seg_. The phenotypic value of the offspring is *z*_*o*_ = *g*_*o*_ + *e*_*o*_, where *e*_*o*_ is environmental noise, which we assume is normally distributed with mean 0 and variance *V*_e_.

Suppose a round of selection changes the phenotypic variance from *V*_*z*_ to 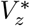. Since we assume both genetic values and environmental noise are normally distributed, by the property of multi-normal distribution, the genetic variance after selection is 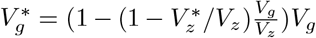. By Equation (S15), the genetic variance in the next generation is

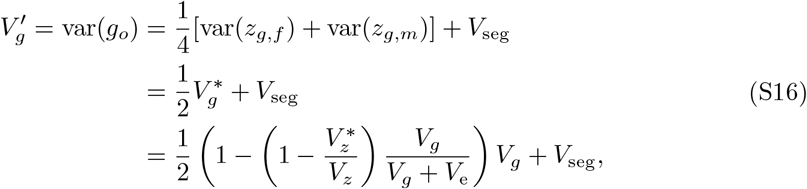

where we assume viability selection acts identically on females and males. The equilibrium genetic variance, 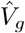, can be obtained by solving this recursion, giving equilibrium phenotypic variance is 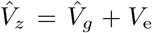. The phenotypic variance after selection, 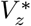 can be calculated based on the distribution of trait values after selection, given by

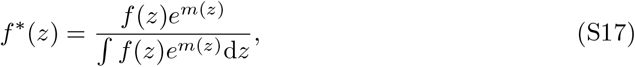

where *f* (*z*) is the trait value distribution before selection. An analytical expression for 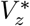 is generally unavailable. However, for the special case when *k* = 2, we have 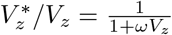. From Equation (S17), the equilibrium genetic variance is then

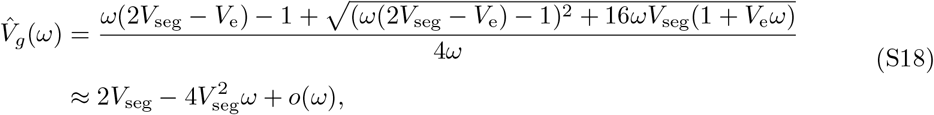

where the approximation assumes weak selection and the first term is the equilibrium value under no selection (Barton et al., 2017).

Figure S1A compares the steady-state fitness predicted by the model incorporating the reduction of genetic variance by selection with the model assuming constant genetic variance (as in the main text). In general, the difference is very slight. This is partly because the reduction of genetic variance has opposing effects on fitness; it reduces the variance load but increases the lag load (Figure S1B), and these two effects partly cancel each other out.

**Figure S1:**
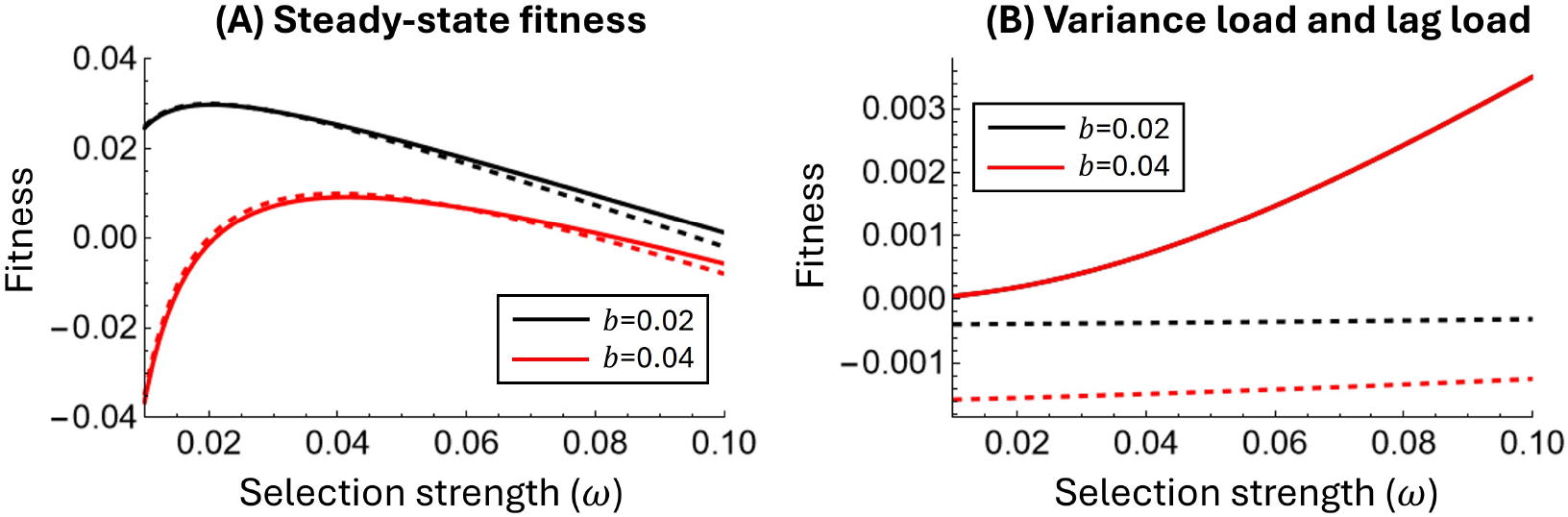
(A) Comparison of predicted steady-state fitness under gradual environmental change. Solid lines: model that incorporates the reduction of genetic variance by selection (Equation (S18)). Dashed lines: model that assumes constant genetic variance (*V*_*g*_ = 2*V*_seg_). (B)Difference in the steady-state fitness reduction contributed by variance load (solid lines) and lag load (dashed lines) between the model with reduced genetic variance and the model with constant genetic variance. The solid black line and solid red line (with *k* = 2 the variance load doesn’t depend on the lag and therefore on *b*). Other parameters: *k* = 2, *m*_0_ = 0.05, *V*_seg_ = 0.5, *V*_e_ = 0.

### S5 Simulations

#### S5.1 Rescue via evolution at a single locus

For rescue via evolution at a single locus, we simulate a population with discrete generations. We assume the offspring number of each individual is Poisson distributed with mean equal to their absolute fitness. Instead of determining the offspring number of each individual according to their absolute fitness, *W*_*i*_, we speed up the simulation by obtaining the total offspring number first (Roze, 2015). Specifically, since Poisson distributions are additive, we first determine the population size *N*_*t*+1_ by drawing from a Poisson distribution with mean equal to the expected total offspring number, 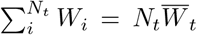. We then calculate the expected genotype frequencies after viability selection, reproduction, and mutation, from which the number of each genotype in the next generation is sampled to make the *N*_*t*+1_ offspring. For rescue from new mutation we expose *N*_0_ individuals with *aa* genotypes to the environmental change. For rescue from standing variation we assume the population size is constant at *N*_0_ before the environmental change and we run the system for 6*N*_0_ generations to reach mutation-selection-drift balance. The simulation stops either when the population goes extinct (*N*_*t*_ = 0) or when the population is considered established 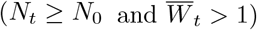.

#### S5.2 Rescue via the evolution of a quantitative trait

For rescue via the evolution of a quantitative trait the simulation considers a random mating, sexually reproducing population with non-overlapping generations. We simulate the evolution of the quantitative trait by using the infinitesimal model (Barton et al., 2017), as described by Equation (S15).

As in the one-locus case, we assume the offspring number of each individual is Poisson distributed with mean equal to its absolute fitness, 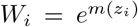. To generate the next generation, we first determine the population size in the next generation by drawing from a Poisson distribution with mean 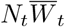. For each offspring, we sample its mother and father from the parental population with probability proportional to their individuals fitnesses.

When there is no selection, the equilibrium genetic variance is 2*V*_seg_. Therefore, to compare the results from simulation with model predictions, we set *V*_seg_ = *V*_*g*_*/*2, where *V*_*g*_ is the genetic variance parameter used in the model. However, in the model, genetic variance is a parameter and assumed to be constant over time. In contrast, in the simulation genetic variance is variable over time, being increased by segregation variance *V*_seg_ and depleted by selection and sexual reproduction each generation. Genetic variance therefore declines with selection.

#### S5.3 Demographic stochasticity

It should be emphasized that when population extinction is due to demographic stochasticity, the strength of demographic stochasticity in our model and that in the simulation slightly differ. Specifically, in the model we assume the variance of population size in the next generation relative to the population size in the current generation is var 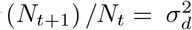, with constant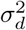. In contrast, in the simulation the variance of population size is var 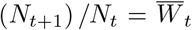, which will therefore change over time. Since we assume weak selection, the strength of demographic stochasticity in the simulation, 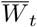, remains close to 1. We therefore use 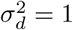 in Figure 1.

## References

Alexander, H. K., Martin, G., Martin, O. Y., & Bonhoeffer, S. (2014). Evolutionary rescue: Linking theory for conservation and medicine. Evolutionary Applications, 7 (10), 1161–1179.

Anciaux, Y., Chevin, L.-M., Ronce, O., & Martin, G. (2018). Evolutionary rescue over a fitness landscape. Genetics, 209 (1), 265–279.

Andersson, D. I., & Hughes, D. (2012). Evolution of antibiotic resistance at non-lethal drug concentrations. Drug Resistance Updates, 15 (3), 162–172.

Barfield, M., Holt, R. D., & Gomulkiewicz, R. (2011). Evolution in stage-structured populations. The American Naturalist, 177 (4), 397–409.

Bell, G. (2017). Evolutionary rescue. Annual Review of Ecology, Evolution, and Systematics, 48, 605–627.

Blanquart, F. (2019). Evolutionary epidemiology models to predict the dynamics of antibiotic resistance. Evolutionary Applications, 12 (3), 365–383.

Bürger, R., & Lynch, M. (1995). Evolution and extinction in a changing environment: A quantitative-genetic analysis. Evolution, 49 (1), 151–163.

Busi, R., Vila-Aiub, M. M., & Powles, S. B. (2011). Genetic control of a cytochrome p450 metabolism-based herbicide resistance mechanism in lolium rigidum. Heredity, 106 (5), 817–824.

Caballero, A., & Hill, W. G. (1992). Effective size of nonrandom mating populations. Genetics, 130 (4), 909–916.

Carlson, S. M., Cunningham, C. J., & Westley, P. A. (2014). Evolutionary rescue in a changing world. Trends in Ecology & Evolution, 29 (9), 521–530.

Charlesworth, B. (2020). How long does it take to fix a favorable mutation, and why should we care? The American Naturalist, 195 (5), 753–771.

Cotto, O., Sandell, L., Chevin, L.-M., & Ronce, O. (2019). Maladaptive shifts in life history in a changing environment. The American Naturalist, 194 (4), 558–573.

Day, T., Huijben, S., & Read, A. F. (2015). Is selection relevant in the evolutionary emergence of drug resistance? Trends in Microbiology, 23 (3), 126–133.

Day, T., & Read, A. F. (2016). Does high-dose antimicrobial chemotherapy prevent the evolution of resistance? PLoS Computational Biology, 12 (1), e1004689.

Glémin, S., & Ronfort, J. (2013). Adaptation and maladaptation in selfing and outcrossing species: New mutations versus standing variation. Evolution, 67 (1), 225–240.

Gomulkiewicz, R., & Holt, R. D. (1995). When does evolution by natural selection prevent extinction? Evolution, 201–207.

Gomulkiewicz, R., Holt, R. D., Barfield, M., & Nuismer, S. L. (2010). Genetics, adaptation, and invasion in harsh environments. Evolutionary Applications, 3 (2), 97–108.

Gould, F., Brown, Z. S., & Kuzma, J. (2018). Wicked evolution: Can we address the sociobiological dilemma of pesticide resistance? Science, 360 (6390), 728–732.

Gullberg, E., Cao, S., Berg, O. G., Ilbäck, C., Sandegren, L., Hughes, D., & Andersson, D. I. (2011). Selection of resistant bacteria at very low antibiotic concentrations. PLoS Pathogens, 7 (7), e1002158.

Haldane, J. B. S. (1927). A mathematical theory of natural and artificial selection, part v: Selection and mutation. Mathematical proceedings of the Cambridge philosophical society, 23 (7), 838–844.

Hartfield, M., & Glémin, S. (2016). Limits to adaptation in partially selfing species. Genetics, 203 (2), 959–974.

Huijben, S., Sim, D. G., Nelson, W. A., & Read, A. F. (2011). The fitness of drug-resistant malaria parasites in a rodent model: Multiplicity of infection. Journal of Evolutionary Biology, 24 (11), 2410–2422.

Kopp, M., & Matuszewski, S. (2014). Rapid evolution of quantitative traits: Theoretical perspectives. Evolutionary Applications, 7 (1), 169–191.

Lande, R. (1976). Natural selection and random genetic drift in phenotypic evolution. Evolution, 314–334.

Lande, R., & Shannon, S. (1996). The role of genetic variation in adaptation and population persistence in a changing environment. Evolution, 434–437.

Lipsitch, M., & Levin, B. R. (1997). The population dynamics of antimicrobial chemotherapy. Antimicrobial Agents and Chemotherapy, 41 (2), 363–373.

Lynch, M., & Lande, R. (1993). Evolution and extinction in response to environmental change. In P. Kareiva, J. G. Kingsolver, & R. B. Huey (Eds.), Biotic interactions and global change (pp. 234–250). Sinauer.

Orr, H. A., & Unckless, R. L. (2008). Population extinction and the genetics of adaptation. The American Naturalist, 172 (2), 160–169.

Osmond, M. M., & Klausmeier, C. A. (2017). An evolutionary tipping point in a changing environment. Evolution, 71 (12), 2930–2941.

Uecker, H., & Hermisson, J. (2016). The role of recombination in evolutionary rescue. Genetics, 202 (2), 721–732.

Uecker, H., Otto, S. P., & Hermisson, J. (2014). Evolutionary rescue in structured popula-tions. The American Naturalist, 183 (1), E17–E35.

Uecker, H. (2017). Evolutionary rescue in randomly mating, selfing, and clonal populations. Evolution, 71 (4), 845–858.

Xu, K., Vision, T. J., & Servedio, M. R. (2023). Evolutionary rescue under demographic and environmental stochasticity. Journal of Evolutionary Biology, 36 (10), 1525–1538.

Zur Wiesch, P. A., Kouyos, R., Engelstädter, J., Regoes, R. R., & Bonhoeffer, S. (2011). Population biological principles of drug-resistance evolution in infectious diseases. The Lancet Infectious Diseases, 11 (3), 236–247.

## References

Barton, N. H., Etheridge, A. M., & Véber, A. (2017). The infinitesimal model: Definition, derivation, and implications. Theoretical population biology, 118, 50–73.

Iwasa, Y., Pomiankowski, A., & Nee, S. (1991). The evolution of costly mate preferences ii. the “handicap” principle. Evolution, 45 (6), 1431–1442.

Price, G. (1970). Selection and covariance. Nature, 227 (5257), 520–521.

Roze, D. (2015). Effects of interference between selected loci on the mutation load, inbreeding depression, and heterosis. Genetics, 201 (2), 745–757.

